# Vertical transmission and community turnover in seed-borne microbiomes: lessons from an aMAIZEing model system

**DOI:** 10.64898/2026.11.16.766985

**Authors:** Sarah Henaut-Jacobs, Sara Sangi, João Pedro Matos, Rafael Chaves Ribeiro, Gabriela Petroceli-Mota, Bruno da Costa Rodrigues, Valter Jário de Lima, Samuel Henrique Kamphorst, Uéliton Alves de Oliveira, Rodrigo Nunes-da-Fonseca, Fabio Lopes Olivares, Rampal S. Etienne, Thiago M. Venancio

**Affiliations:** Laboratório de Química e Função de Proteínas e Peptídeos (LQFPP), Biology and Biotechnology Center, State University of Northern Rio de Janeiro, Campos dos Goytacazes, Brazil; Groningen Institute for Evolutionary Life Sciences, University of Groningen, Groningen, The Netherlands; Laboratório de Biologia Celular e Tecidual (LBCT), Biology and Biotechnology Center, State University of Northern Rio de Janeiro, Campos dos Goytacazes, Brazil; Instituto de Biodiversidade e Sustentabilidade (NUPEM/UFRJ), Federal University of Rio de Janeiro, Macaé, RJ, Brazil, Avenida São José do Barreto, Macaé, RJ, 27965-045, Brazil; Laboratory of Plant Genetics and Breeding, Agricultural Science and Technology Center, State University of Northern Rio de Janeiro, Campos dos Goytacazes, RJ, 28013-602, Brazil; Center for Agricultural and Biological Sciences, Vale do Acaraú State University – Campus Ibiapaba, São Benedito, CE, 62370-000, Brazil; Latin American Institute of Life and Natural Sciences, Federal University of Latin American Integration, Foz do Iguaçu, PR, 85867-970, Brazil

**Author notes:** Corresponding authors: SH-J; TMV.

## Abstract

The seed microbiome represents a specialized ecological niche, governed by assembly processes that are only partially coupled to the surrounding soil community. While environmental recruitment and vertical transmission from the maternal plant are both recognized sources of seed-associated microorganisms, the mechanisms that determine their relative contributions and whether these contributions shift across generations remain poorly understood. Here, we used long-read metagenomic sequencing to track seed-borne bacterial community dynamics across a founding seed pool and two subsequent generations in two divergent maize (*Zea mays* L.) varieties exposed to contrasting organic and mineral fertilisation regimes. We recovered 112 medium- or high-quality metagenome-assembled genomes and combined MAG-level taxonomic and functional analyses with genus-level community profiling to characterize persistence and turnover dynamics within the seed microbiome. While soil taxonomic diversity was high, seed-associated communities showed a near-complete compositional disconnect from soil-dominant assemblages taxa, with a persistent Firmicutes core maintained across generations, consistent with vertical transmission. Priority effects remain a possible explanation for these patterns but were not directly tested. Genus-level source attribution suggested greater environmental recruitment under organic fertilization, with more diverse soil communities and a stronger contribution from the preceding seed generation under mineral fertilization. Seed-associated MAGs also carried genes associated with plant growth-promoting potential, although these detections could not be consistently linked to specific transmission routes. In contrast, maize genotype had no detectable effect on seed microbiome composition. Intergenerational persistence and environmental recruitment jointly shape seed-derived communities under contrasting fertilization regimes. Their relative contributions remain inferred from taxonomic overlap, and the effects of microbial source-pool diversity cannot be separated from the accompanying differences in soil chemistry in this design.

## INTRODUCTION

The plant microbiome is assembled from two primary sources: recruitment from the surrounding soil and vertical transmission from the mother plant (the seed-borne microbiome) ^1,2^. While the soil is recognized as the primary reservoir for plant-associated microbes ^3,4^, the seed-borne community represents a specialized microbial niche that is only partly coupled to the surrounding soil community and remains poorly characterized. This inherited microbiome plays a critical role in plant development, providing immediate protection against pathogens and promoting seedling growth ^5,6^. Furthermore, micro-organisms require specific functional traits and colonization mechanisms to establish within the seed environment ^7^, suggesting that assembly is not purely stochastic.

If assembly is structured rather than random, the question becomes which processes do the structuring. Viewing the seed as a microbial community partly decoupled from the surrounding soil brings its own species interactions and environmental pressures into focus. We previously observed compositional divergences among maize varieties, consistent with host genotype acting as a selective filter on community assembly ^8^. More broadly, community assembly in the phytobiome is best understood within a metacommunity framework, where local community composition reflects the interplay between dispersal from the regional species pool, environmental filtering, and stochastic colonization events ^9,10^. From this perspective, host genotype constitutes one filter among several: the structure of the surrounding soil community determines which colonizers are available, while abiotic gradients, priority effects, and niche preemption further shape which taxa successfully establish within the seed ^11–13^. Disentangling these overlapping drivers is therefore essential for understanding what governs the composition of the inherited microbiome.

These assembly dynamics have implications beyond a single generation. Seedlings growing in soils with a low abundance of potential colonizers are more likely to be re-colonized by members of the previous generation’s microbiome ^14^, suggesting that vertical transmission through the seed can function as a buffer against environmental microbial scarcity. Whether this buffering compounds across successive generations, however, has not been tested, as existing evidence is based on single-generation observations. This points toward a broader evolutionary question: whether the selective retention of specific taxa across generations represents an adaptive strategy, analogous to heritable symbiont dynamics described in animal systems ^15,16^, even though seed transmission lacks the obligate dependence and dedicated cellular machinery characteristic of insect endosymbioses. In such a scenario, the host maintains a core microbiome that confers fitness benefits. Such benefits may operate across multiple dimensions, enhancing germination success, competitive establishment under resource limitation, or resistance to abiotic and biotic stress, but the degree to which the inherited community shapes these outcomes remains unresolved. Critically, it is also unclear whether this transgenerational continuity persists under strong environmental perturbation or breaks down when external selective pressures overwhelm the vertical transmission signal. These open questions motivate a longitudinal, multi-generational perspective on seed microbiome dynamics.

Intensive agriculture imposes strong selective pressures on soil ecosystems, altering natural biogeochemical cycles and belowground community structure. Environmental homogenization typical of intensive farming drives substantial depletion of soil biodiversity across multiple trophic levels, simplifying microbial networks and reducing the diversity of the coloniser pool available to plants ^17^. If seed microbiome assembly depends on dispersal from that pool, agricultural intensification should shift the balance between environmental recruitment and vertical transmission, a prediction that has not been tested across generations.

A primary driver of this ecological degradation is the shift in carbon input dynamics. In natural, less disturbed systems, the availability of complex, diverse biological carbon sources, such as the products of decomposition, enforces niche partitioning. This sustains high metabolic diversity and complex cross-feeding networks among microbes ^18^. Conversely, modern agricultural regimes often supply the system with high loads of synthetic chemical inputs. These inputs supply mineral nutrients without the structurally complex organic carbon that sustains microbial metabolic diversity, even where total soil organic matter is comparable. The resulting abundance of readily accessible nutrients may relax the selective pressures that favor metabolic cooperation. Consistent with the Black Queen Hypothesis, this constant exogenous supply of resources can drive reductive evolution, wherein community members streamline their genomes by shedding the energetically expensive genes required to process complex organic matter ^19^. Ultimately, substituting complex biological carbon with simplified synthetic alternatives reduces the structural complexity of the soil microbiome, limiting its adaptive capacity and resilience to further environmental stress ^20,21^.

In this work, we employed maize as a model system to investigate transgenerational shifts in the seed-borne microbiome under contrasting organic and mineral fertilization regimes that serve as tractable proxies for the carbon-input dynamics of low-input versus intensive agricultural systems. To isolate the effects of environmental selection from host genetics, we ran the experiment in parallel across two divergent maize varieties. Using long-read sequencing, we then tracked microbial community dynamics within the seeds across three consecutive generations, recovering metagenome-assembled genomes (MAGs) and mapping functional trajectories of the microbiome over time.

## MATERIALS AND METHODS

### Experimental design

Two maize (*Zea mays* L.) varieties were selected from the Anchieta diversity hotspot in Santa Catarina, Brazil, due to their shared geographical origins and biological diversity. The first variety, commercial BNS Bandeirantes-Bonamigo (MOD), is a rustic commercial variety launched in 2001, bred for grain production (Sementes Bonamigo, 2024). The second variety is a semi-domesticated landrace (LAND) maintained by local smallholder farmers in the same region. The material was collected and registered under SisGen (Sistema Nacional de Gestão do Patrimônio Genético e do Conhecimento Tradicional Associado) record number 83527850015.

We used a soil classified as a Latossolo under the Brazilian Soil Classification System (SiBCS), collected in northern Rio de Janeiro, Brazil, as the experimental substrate. The soil was sifted and homogenised in batches of 500kg using a large container, and its pH was adjusted with liming to ∼6.5 for maize production. Soil micronutrient levels and pH were characterized through biochemical analysis (Table 1) (FUNDENOR - Campos dos Goytacazes, Brazil).

**Table 1:** Physicochemical characteristics of the experimental soils. Soil samples (*n* = 3 replicates per treatment) were collected from the surface level of the tubes before planting. Chemical analyses were performed by FUNDENOR (Campos dos Goytacazes, Brazil).

| **Parameter** | **Description** | **Bio treatment Averages** | **Chem treatment Averages** |
|---|---|---|---|
| **pH** | Soil Acidity | 5.4 ± 0.0 | 4.5 ± 0.17 |
| **Al (mmolc/dm³)** | Exchangeable Aluminum | 0.4 ± 0.1 | 6.53 ± 1.78 |
| **Al S (%)** | Aluminum Saturation | 1 ± 0.0 | 11.33 ± 3.79 |
| **BS (%)** | Base Saturation | 73.6 ± 2.12 | 55 ± 3.61 |
| **P (mg/dm³)** | Phosphorus | 116 ± 7.55 | 75 ± 22.61 |
| **SOM (g/dm³)** | Soil Organic Matter | 20.8 ± 1.71 | 17.8 ± 0.92 |

Two contrasting fertilizer regimes were simulated in a controlled greenhouse setting: a mineral chemical farming treatment and an organic fertilizer treatment. The inorganic chemical farming treatment was applied using phosphorus (P₂O₅) at 145 mg/kg, urea at 40 mg/kg, and potassium chloride at 34 mg/kg, representing a conventional synthetic fertilization regime (hereafter chemical, c). The organic treatment consisted solely of vermicompost at 20 g/kg, providing a complex organic carbon source instead of synthetic inputs (hereafter biological, b). As a consequence of the fertilization regimes, the treatments diverged in soil pH and exchangeable aluminum (Table 1). Treatment contrasts should therefore be interpreted as composite differences between input regimes rather than as isolated effects of nutrient source. Top-dressing fertilization was applied at 15 and 30 days post-planting. The chemical treatment received a slow-release NPK fertilizer, while the biological treatment received a combination of castor bean cake and bone meal. Each treatment-variety combination was planted in eight separate tubes, staggered in pairs at one-week intervals to ensure simultaneous availability of pollen and receptive flowers for cross-pollination. Plants were grown in PVC tubes containing 15 kg of treated soil per tube to ensure standardized rooting volume and minimize inter-replicate soil variability.

To produce the second generation (g2), seeds harvested from the first generation (g1) were pooled by treatment-variety combination and planted in fresh soil prepared with the identical fertilization regimes described above.

### Pollination

Controlled pollination was conducted through crosses among distinct parental plants within each variety, avoiding inbreeding, with parental plants tracked throughout the process. To avoid cross-contamination, maize ears were covered before silk appeared, and pollinated corn silks were immediately covered with paper bags until fertilization was complete. Upon maturation, cobs were harvested and dried at 28-35°C for seven days to preserve seed viability. Seeds were subsequently stored at 5°C between experiments to preserve their viability.

### Generations

Post-harvest, all seeds produced were pooled, and subsets were randomly assigned to either microbiome extraction or the growing of the next generation. This longitudinal tracking encompassed three generations: a founding seed pool (g0) prior to treatment, followed by two generations (g1 and g2) grown under biological (b) and chemical (c) fertilization regimes, enabling a multi-generational assessment of community trajectories. Within each treatment arm, every generation was sown from seed harvested from the preceding generation of the same arm (g0→g1→g2), ensuring that the biological and chemical lineages were propagated independently across generations (Figure 1). Samples are labeled by treatment prefix and generation (e.g., bg1 = biological, generation 1; cg2 = chemical, generation 2). Three replicates were taken out of each pool of seeds and denoted as _1, _2, and _3.

**Figure 1:**
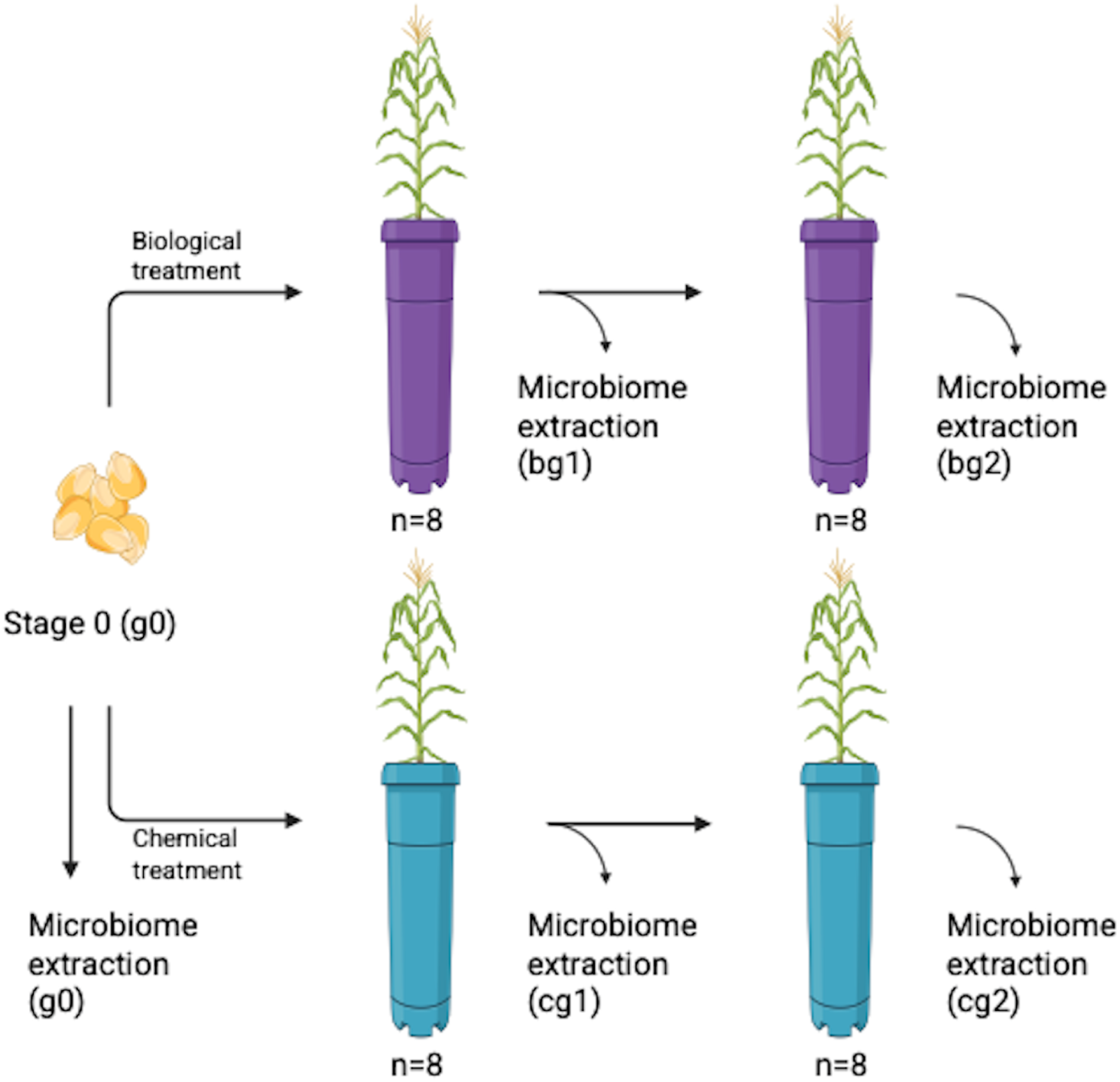
Experimental design across the founding seed pool (g0) of the genotypes and two subsequent generations (g1, g2) propagated independently under biological (b) and chemical (c) treatments. Plants in the subsequent generations were planted from a pool of seeds from the previous generation. This procedure was repeated for both genotypes used in this research

### Seed-Borne Microbiome Extraction

Seed-borne microbiome isolation was initiated within 10 days post-maturation to minimize community drift during storage. For each of the three replicates, 30 maize seeds were placed in a 3 L hermetically sealed glass jar containing 750 g of sterilized Basaplant substrate and 100 mL of sterilized water, and incubated in a BOD chamber at 28°C for seven days under a 12-hour photoperiod. This germination-based approach allows seed-borne microbes to colonize the emerging rhizosphere, providing sufficient biomass for downstream DNA extraction. Because seed-borne taxa exert their effects on the host through germination and colonization of the emerging seedling, this approach captures the functionally realized fraction of the seed-associated community (i.e., the organisms that successfully establish) rather than the total pool of seed-resident DNA, which includes dormant and non-viable cells. A separate jar containing only the sterilized Basaplant with sterilized water was also placed in the BOD to serve as the negative control for the media used in the following steps. Following incubation, rhizosphere material was collected, and DNA was extracted using the Qiagen DNeasy PowerSoil kit following the manufacturer’s protocol. Extracted DNA was stored at -80°C until sequencing. Microbiome analyses were additionally performed on untreated seeds before any experimental generation (stage 0) using the same protocol, providing a baseline community profile. Soil samples from both treatment groups were also collected at each generation and sequenced to characterize the pool of potential microbial colonizers available to each cohort. To ensure high-quality metagenomic data, genomic material was quantified via spectrophotometry, and only those extractions exceeding 20 ng/µl were prioritized for downstream sequencing. Seeds harvested from the eight replicate tubes of each treatment-variety combination were combined and distributed into three germination jars, each processed as an independent extraction replicate.

### Sequencing

Libraries were prepared using the SQK-RBK114-96 Rapid Barcoding Kit (Oxford Nanopore Technologies) according to the manufacturer’s instructions and sequenced on a PromethION 2 Solo platform. The complete pool of barcoded libraries was sequenced across two independent flow cells, with sequencing runs maintained until pore activity was exhausted to maximize total data yield. This strategy generated long-read metagenomic data for all samples. Sequencing yield and per-sample read statistics are provided in Table S1.

### Data cleaning

Basecalling, adapter trimming, and demultiplexing were performed using Dorado (v1.2.0) (https://github.com/nanoporetech/dorado). Reads shorter than 500 bp or with a mean quality score below Q9 were discarded using NanoFilt (v2.8.0) ^22^. Host decontamination was carried out with Hostile (v2.0.20) ^23^, removing reads aligning to the maize reference genome (GCF_902167145.1). Reads identified in negative control samples were treated as exogenous DNA contaminants and removed using Hostile with a minimum minimap2 v(2.30)^24^ chaining score of 150 (-m 150), a more stringent threshold than Hostile’s default of 40, applied to minimise false positive removal of genuine microbial reads while ensuring only confidently aligned contaminant sequences were excluded from downstream analyses.

### Assembly and binning

Metagenomic assembly was performed using Flye (v2.9.6) ^25^. Reads were mapped back to the assembly using Minimap2 (v2.30) ^24^ to generate coverage information for binning. Genomic bins were constructed with MetaBAT2 (v2.18) ^26^ and evaluated for completeness and contamination using CheckM2 (v1.1.0) ^27^. Bins were classified as high-, medium-, or low-quality metagenome-assembled genomes (MAGs) according to the MIMAG standards ^28^. Taxonomic classification of MAGs was performed using GTDB-Tk (v2.5.2 release 226) ^29^, and broader contig-level taxonomic assignment was carried out with Kraken2 (v2.1.6 plus PFP 20250714) ^30^. Because some individual libraries from these low-biomass samples did not reach the coverage required for reliable binning, reads from the three replicates of each treatment were pooled into a single file prior to assembly, enabling MAG recovery and finer taxonomic resolution (Table S2). As a consequence, MAG-level taxonomy results represent a single assembly per treatment and are presented descriptively; no within-treatment variance is available at this level, and MAG-based comparisons between treatments are not statistically tested. Pooling reads across replicates can affect binning when strain-level variation is present; contamination estimates from CheckM2 were used to identify and exclude affected bins, and only medium- and high-quality MAGs that met MIMAG standards were retained.

### Plant growth promotion and sporulation traits

To explore specific genomic capacities of target taxa, we used individual replicate-assembled MAGs.

An *in-house* database for known plant growth promotion-associated genes (PGP-associated genes) ^31^ was used to infer plant growth promotion potential. A local database containing genes associated with sporulation capacity was created based on the work of Galperin et al. (2022) ^32^. We used Usearch (v11.0.667_i86linux32) to align MAGs against both databases, using identity thresholds of 50% and 60% coverage between predicted proteins from the MAGs and the protein database ^33^. To double-check the complete absence of certain genes, we conducted the same Usearch analysis on unbinned FASTA files.

### General taxonomic profile statistics

Bray-Curtis dissimilarities were calculated using the Vegan package (v2.7.2) ^34^ in R (v4.5.0). To visualize differences in community composition, Principal Coordinates Analysis (PCoA) was performed on the Bray-Curtis matrix using the ape package (v5.8-1) ^35^. Statistical differences in community structure among samples were evaluated using Permutational Multivariate Analysis of Variance (PERMANOVA) via the adonis2 function in Vegan. The PERMANOVA was conducted using 1000 permutations, strictly constrained within treatment lineages (strata) to account for the non-independence of the generational experimental design.

To explore the trends in the influence of soil community composition on environmental recruitment into the seed microbiome, a beta regression model was fitted for each bacterial class detected in soil using the betareg package (v3.2-2) ^36^ in R, following the methodology from Walsh et al. (2021) ^14^. The predictor was the relative abundance of each bacterial class in the corresponding soil sample, calculated from Kraken2 class-level counts normalized within each soil sample. This was calculated from Kraken2 class-level counts and normalized within each soil sample. The unit of observation was individual soil samples (bg1, bg2, cg1, cg2) matched to their corresponding seed community. Models were fitted only for classes with more than two matched observations, and p-values were corrected for multiple testing using the Benjamini-Hochberg procedure. Nine classes were excluded from the final figure as their slopes were estimated over insufficient predictor variance to be interpretable (showing up in less than three replicates).

To visualise the relative contributions of vertical transmission and environmental recruitment to seed community assembly across generations, genus-level source attribution was performed for each generation transition using predefined source-destination pairs: Stage 0 seeds to g1 seeds, soil samples to g1 seeds, g1 seeds to g2 seeds, and soil samples to g2 seeds. Attribution was based on genus presence/absence: a genus detected in a destination sample was attributed to a given source if it was also detected in that source sample, with a minimum detection threshold of ≥2 reads. Genera present in the destination but absent from all defined source pools were classified as unknown origin, reflecting taxa potentially introduced from uncontrolled environmental sources inherent to greenhouse conditions. Genera detected in both the prior seed generation and the corresponding soil sample were attributed to both sources (previous generation and previous soil microbiomes) simultaneously, as the data do not permit unambiguous resolution of their proximate origin. This additive attribution scheme means that proportions reported for soil recruitment and vertical transmission are not mutually exclusive for shared taxa, and the reported percentages should be interpreted as upper bounds on each source’s contribution rather than strict partitions. All flows were quantified as the number of genera attributed to each source rather than read abundance, giving equal weight to each taxon regardless of its relative abundance in the community. To assess the sensitivity of the reported contrasts to this attribution choice, we additionally computed proportions under two conservative scenarios: assigning all shared taxa exclusively to the prior seed (minimising the soil contribution) and exclusively to soil (minimising the vertical contribution). Under both alternative schemes the directional contrast between biological and chemical treatment lineages was preserved, with biological treatments consistently showing higher soil recruitment than chemical treatments regardless of how shared taxa were handled.

## RESULTS AND DISCUSSION

### Transgenerational shifts in seed-borne community composition: persistent and displaced taxa

Community composition of the seed-borne microbiome differed significantly across treatment-variety groups. PERMANOVA analysis, conducted on seed samples only with soil samples excluded to prevent their disproportionate influence on treatment-level effects, revealed that treatment identity explained 57% of the total variance in community composition (R² = 0.57, p < 0.002; 1000 permutations constrained by treatment lineage), furthermore betadisper tests confirmed that this result reflects true centroid differences rather than dispersion artefacts (p > 0.05). Ordination along the first two principal coordinates captured 31.4% of total variance (Figure S1) (PCo1 = 21.8%, PCo2 = 9.7%), with samples separating along PCo1 broadly according to treatment type and along PCo2 according to host genotype, suggesting independent but concurrent effects of environmental filtering and host-mediated selection on community structure.

Genus-level profiles revealed a marked difference between the dominant taxa in soil and those recovered from seed-derived communities (Figure 2). The top ten genera in soil collectively accounted for 64% of all soil reads yet contributed only 2.3% of seed community reads across all treatment generations; *Streptomyces* and *Bradyrhizobium* alone represented 32.3% and 17.3% of soil reads, respectively, and were almost absent in seed reads (<0.6%). Conversely, 31% of seed-associated genera had no representation in soil whatsoever, while the remaining 69% were present in soil at abundances too low to translate into meaningful seed colonisation, thus, dominance in the soil community did not translate directly into dominance in the communities recovered after germination. This asymmetry is also visible in the replicate-level overlap shown in Figure 2. Soil replicates shared only a small fraction of their detected genera, with many genera restricted to a single replicate (up to 401 in cg1 and 307 in bg1), a pattern that is expected from soil samples due to their heterogeneous surface and composition, and considerable variation in the size of the shared core between soil samples (7 to 293 genera). This pattern is expected for a highly rich community in which most of the detected diversity consists of rare taxa contributing negligibly to total abundance, as reflected in the small cumulative share of the "Other" category relative to the number of genera it contains. Spatial heterogeneity and sampling variation may also have contributed; replicate-exclusive detections should therefore not be attributed to a single process. Despite *Streptomyces* and *Bradyrhizobium* collectively dominating bacterial abundance in both soil types, neither genus was detected as a meaningful component in any seed-associated sample in any generation. Because DNA was extracted from the rhizosphere of emerging seedlings, the recovered profiles represent organisms that established under the assay conditions rather than the entire seed-resident community. Two non-mutually exclusive mechanisms may explain this disconnect. The first is a colonization competence filter: successful seed entry may require specific traits that many abundant soil taxa simply do not possess, such as surface adhesin expression, tolerance of low water activity, resistance to seed antimicrobial compounds, and cell invasion machinery ^37,38^. The second is a priority effect operating through the existing seed community: taxa already established within the maternal seed microbiome occupy available niches ahead of arriving soil recruits, suppressing their establishment through competitive exclusion regardless of their abundance in the surrounding environment ^39,40^. In this case, however, we would be able to observe non-random co-occurrence of certain taxa, which is not always consistent with our observations (i.e., almost complete absence of the very abundant genus *Burkholderia* after g0, this shows that, despite initial abundance, early occurrence does not guarantee persistence). With compositional data alone, these mechanisms cannot be disentangled; our results are consistent with either mechanism operating alone or both operating concurrently, and distinguishing between them would require targeted approaches such as functional trait profiling or colonization assays.

**Figure 2:**
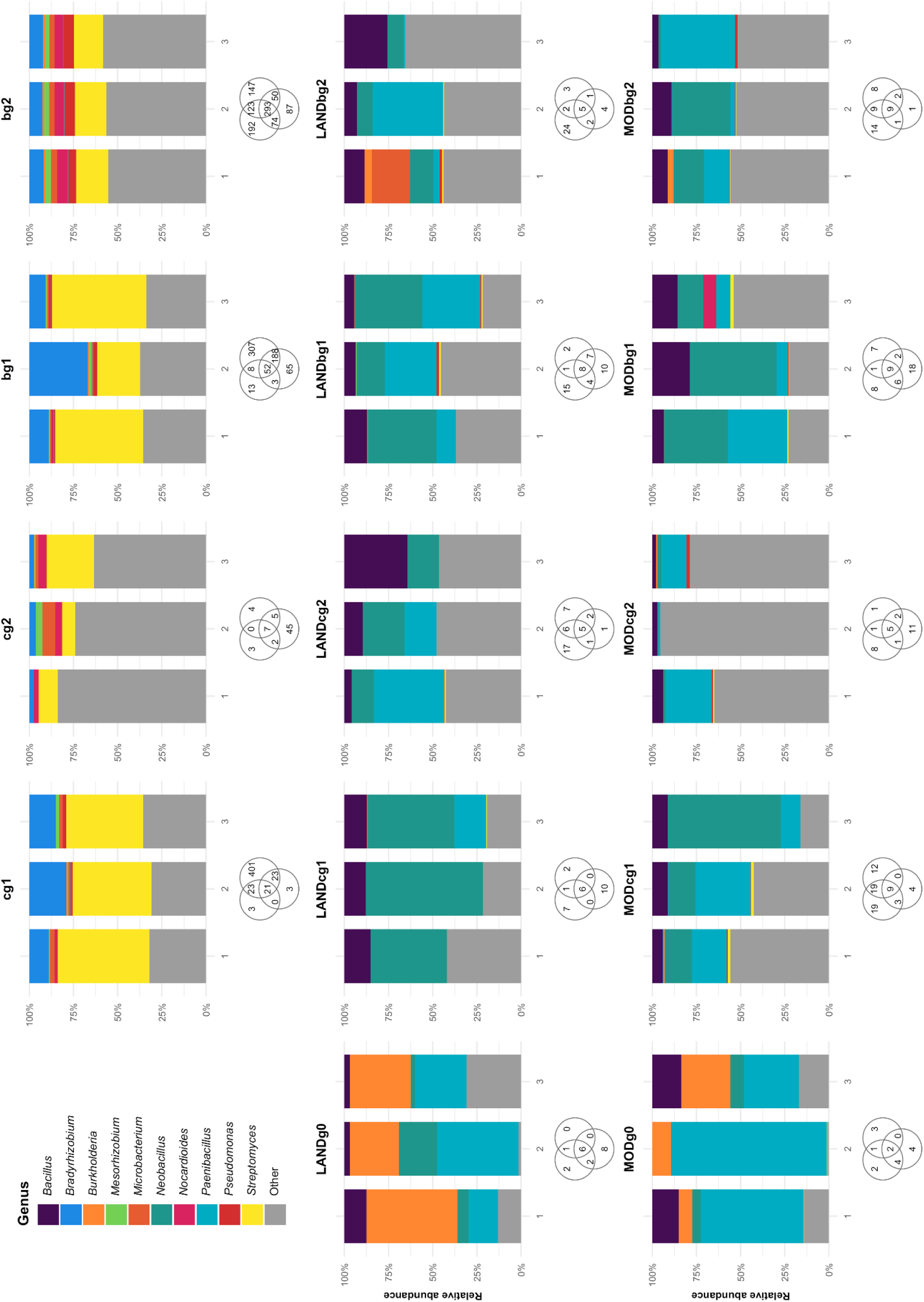
Relative abundance of the ten most abundant bacterial genera in each sample group (n = 3 replicates per panel). Panel labels indicate sample type and experimental stage: bulk soil samples (cg1, cg2, bg1, and bg2) are denoted by treatment-generation, initial baseline seeds (LANDg0 and MODg0) by Variety-Generation, and all other seed samples by Variety-Treatment-Generation. The Venn diagrams below the bar plots depict the overlap of identified genera across the three replicates. All genera outside the top 10 are pooled under “Other”.

Apart from the disconnection between soil and seed microbiomes, the dynamics of specific taxa across generations reveal a biologically interpretable pattern of persistence and turnover. In this study, we use the term "persistent taxa" to describe genera that are recovered consistently across multiple generations within a treatment lineage. Conversely, we use displaced taxa to refer to those that were initially prominent but are subsequently lost or reduced to trace abundances in later generations. Among the persistent taxa, *Paenibacillus*, *Neobacillus,* and *Bacillus* stand out as the most stable components of the seed-borne microbiome: all three were present and prevalent in Stage 0 seeds and maintained across g1 and g2 samples regardless of treatment type or host variety. Seed samples showed a markedly different overlap structure compared with soil: each seed panel converged on a small core of genera shared by all three replicates (5 to 9 genera in each seed sample, including both g0 baselines), and this core consistently comprised the abundant Firmicutes taxa described above. Replicate-exclusive genera in seeds were far fewer than in soil and were confined to the rare fraction, indicating that the dominant structure of the seed community is reproducible across biological replicates rather than assembled stochastically at each germination event. Their combined abundance accounted for 71.7% of total fragments in bg1 and 67.3% in cg1 samples. The persistence of this Firmicutes core in the near-complete absence of a soil source constitutes the strongest evidence for vertical transmission in our dataset. Spore formation provides the most parsimonious physiological explanation, as endospore-forming Bacillales are uniquely equipped to survive seed desiccation, storage, and the physicochemical stresses within the seed interior across successive generations ^41,42^ (see the section Genomic evidence for sporulation potential in the persistent Firmicutes core). From a biotechnological perspective, this multi-generational persistence is notable: these genera are essentially absent from bulk soil in our experiment, yet sustain detectable seed populations two generations after their last plausible soil-based recruitment, suggesting that seed inoculation with Bacillales-based consortia could exert durable transgenerational effects without requiring re-application at every cultivation cycle ^43,44^. Such persistence and any associated benefits would, however, require direct validation for the inoculated strains.

However, this Firmicutes core did not maintain its relative dominance indefinitely. Comparing g1 and g2 across both treatment types, the combined relative abundance of the three persistent genera declined from ∼70% to ∼47% of total fragments, a reduction that was mirrored by a visible increase in the contribution of rarer, less abundant genera (the "Other" category) across all treatment groups (Figure 2). This succession dynamic, in which a dominant founding consortium is progressively diluted by an expanding pool of rarer taxa, is consistent with a model of secondary community diversification following an initial colonization bottleneck ^10^. It remains unclear whether this shift represents passive stochastic colonization or active niche opening, as the Firmicutes core modifies the seed microenvironment for secondary colonizers.

The fate of *Burkholderia* stands apart from the general . This genus was among the most abundant taxa at Stage 0 in both varieties, and at the MAG level, the only genera shared between MODg0 and LANDg0 seeds were traceable to *Burkholderia*-related lineages (Figure 5), indicating that this genus was a reliable component of the maternal seed legacy regardless of host genotype. Yet *Burkholderia* was effectively displaced from g1 onwards across all treatment lineages, declining to negligible abundance by g2 despite being present in the soil colonizer pool available for re-recruitment. This is a fundamentally different pattern from the soil-dominant taxa (*Streptomyces*, *Bradyrhizobium*), which were absent throughout: *Burkholderia* was present in the maternal seed, present in low abundances in soil, and yet still displaced. The most ecologically coherent interpretation is competitive exclusion during seedling establishment: *Burkholderia*, a well-documented endophytic seed coloniser and plant growth-promoting associate ^45,46^, may be outcompeted during germination by the spore-forming Firmicutes that consolidate rapidly in the rhizosphere environment, losing the priority advantage it held in the ungerminated seed. Alternatively, the specific carbon substrates and microbial interaction networks that sustain *Burkholderia*’s seed association on the mother plant may not be replicated in experimental soil conditions, pointing to the context-dependence of vertical transmission for taxa that lack spore-based dormancy mechanisms.

Alpha diversity, assessed by the Shannon index across seed-borne communities (with soil samples excluded), remained statistically stable across all treatment lineages and generations (Kruskal-Wallis, H = 7.51, p = 0.584; Figure 3). At Stage 0, the seed-borne communities of both varieties showed intermediate but variable Shannon diversity (H’ = 3.43 ± 0.55), with MODg0 exhibiting notably high within-group variance (σ = 0.83) compared to the landrace (σ = 0.26), suggesting that the commercial variety’s seed microbiome may be more sensitive to stochastic variation in maternal plant conditions ^47,48^. While a single replicate within the MODcg2 lineage initially appeared to exhibit a sharp drop in diversity, sequencing metrics confirmed that this variance was a technical artefact driven by exceptionally low read depth in that specific sample (Table S1), rather than a biological treatment effect. When contextualised with the compositional shifts observed earlier, The observed compositional differences did not result in a statistically detectable difference in Shannon diversity. This supports a distinction between taxonomic turnover and changes in this diversity metric, rather than demonstrating equivalent diversity among treatments.

**Figure 3:**
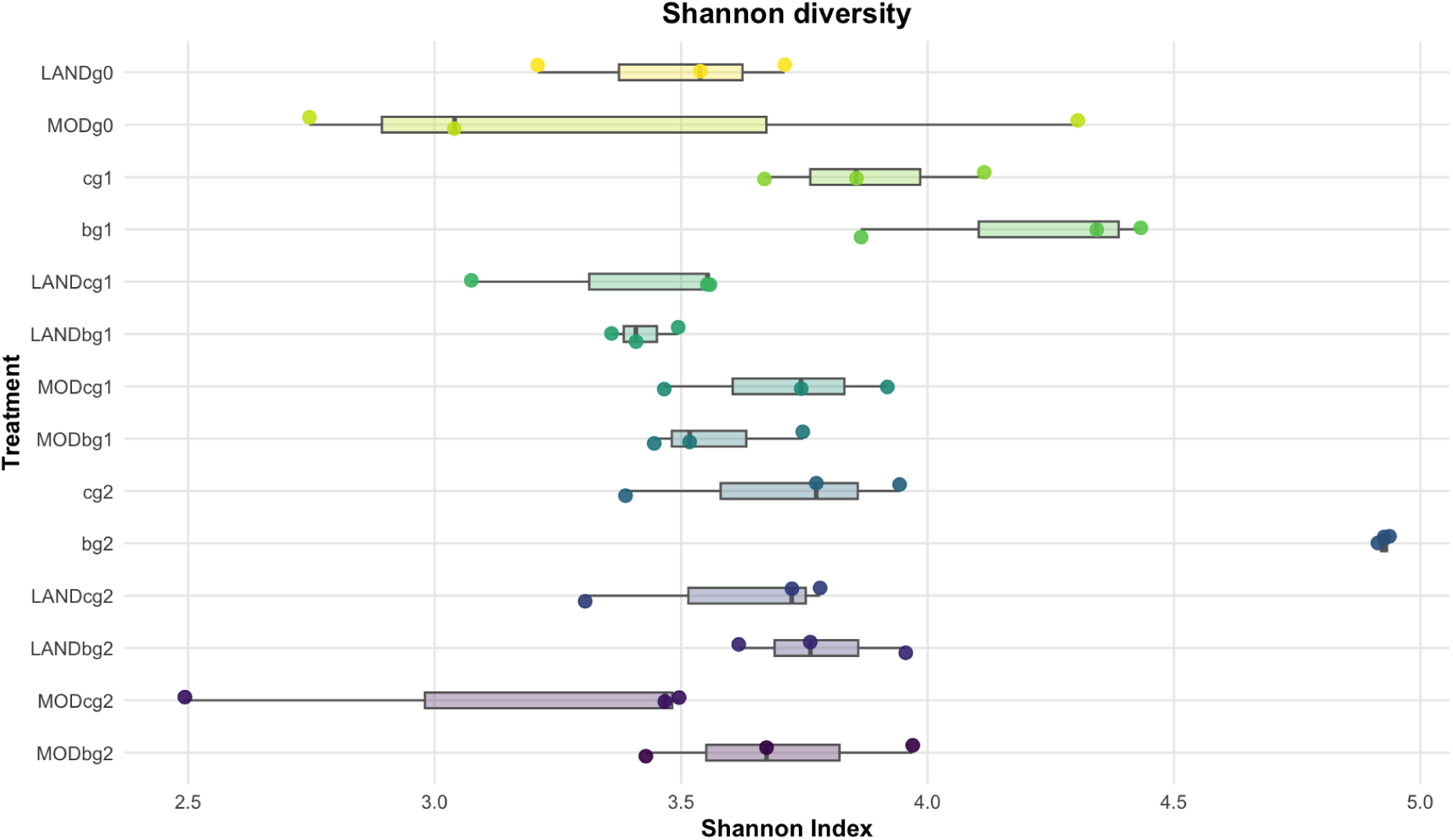
Shannon diversity of soil and seed-derived communities across sample groups. The founding seed groups (LANDg0 and MODg0) appear at the top, followed by groups from subsequent generations. Individual replicate values are shown. The Kruskal-Wallis comparison reported in the text includes seed-derived samples only; soil samples are excluded from that test. The y-axis shows the stage-0 of both seeds on top, followed by next generations.

### Inferred vertical transmission and environmental recruitment

Genus-level transfer analysis (Figures 4 a and b) revealed a consistent but asymmetric contribution of vertical transmission and soil recruitment to community assembly, with the direction of the asymmetry differing fundamentally between treatment types. Before interpreting these proportions, an important caveat applies: a fraction of genera in each generation were assigned to an "unknown" source, meaning they could not be traced to either the prior generation’s seed community or the corresponding soil sample. In a controlled greenhouse setting, complete environmental sterility is unachievable, and a proportion of seed colonisers will inevitably originate from ambient sources (e.g., irrigation water, air deposition, growth setup surfaces, and cross-contamination during handling) that fall outside the tracked pools. Given its substantial magnitude, particularly within the chemical treatment, this “unknown” fraction represents a key gap in the source model. Rather than being non-informative, this block suggests two important non-mutually exclusive possibilities: either the chemical treatment seed communities are more susceptible to colonization from ambient sources, or the tracked seed and soil pools failed to fully capture the diversity of the available microbial reservoirs due to the low abundance of certain taxa in the soil.

**Figure 4.**
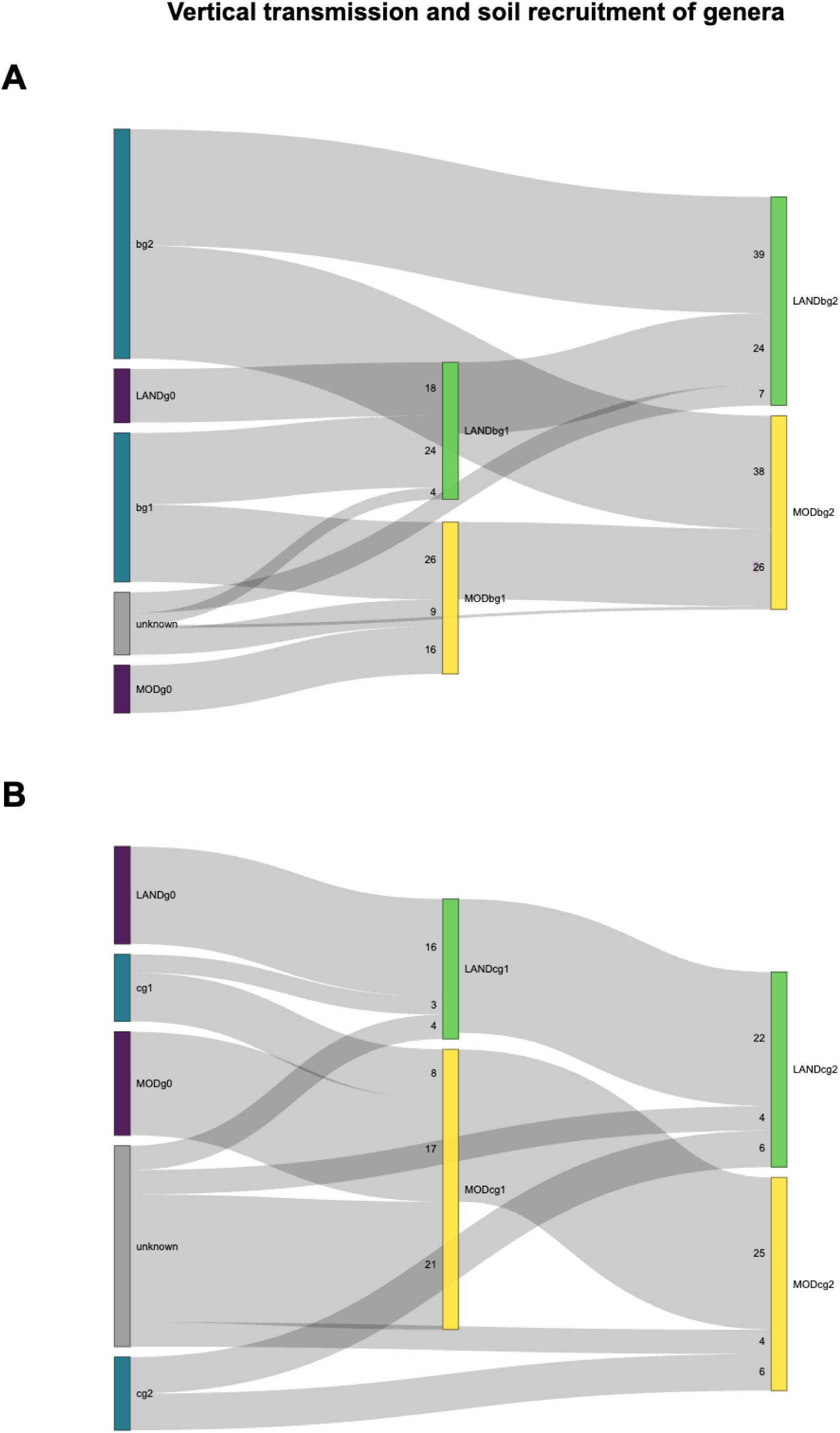
Genus-level source attribution of seed-borne communities across generations for the **(a)** biological and **(b)** chemical carbon source treatments. Flow width is proportional to the number of genera attributed to each source-destination pair. For each generation, the community is partitioned into three sources: genera inherited from the prior seed generation (vertical transmission), genera recruited from the corresponding soil sample (environmental recruitment), and genera absent from all tracked source pools (unknown origin, reflecting uncontrolled environmental inputs inherent to greenhouse conditions). The shifting balance between these three sources across generations reflects the interplay between vertical inheritance and soil-mediated assembly under contrasting agricultural management regimes.

In the b treatment lineages, soil was the dominant source of genera at both g1 and g2, contributing approximately half of all detected genera per generation (51-52% at g1; 56-58% at g2), with the prior seed generation accounting for 31-40% and unknown sources for 2-18% (Figure 4.a). These patterns are consistent with greater environmental recruitment from the more diverse b soil communities. Notably, the unknown fraction declined substantially from g1 (9-18%) to g2 (2-10%). To discriminate whether these unknowns originated from the initial soil inocula or from external environmental sources we compared their identities across the two soil treatments. The unknown taxa were heavily treatment-specific, indicating that they were likely derived from the respective soil communities but fell below the detection threshold of our source-tracking due to their low overall abundance. The decline in their proportions suggests that the system stabilizes, with initially rare taxa being assimilated by the seed-borne microbiome by generation 2, not being flagged as unknown anymore.

The c treatment lineages exhibited a strikingly different assembly dynamic (Figure 4.b). Soil contributed only 13-19% of genera per generation, while the prior seed generation accounted for 37-71% of the community by g2. This progressive dominance of the vertical transmission signal in the chemical lineage is not straightforwardly interpretable as greater host fidelity to the inherited microbiome. Rather, it likely reflects the impoverishment of the c soil coloniser pool documented above: when the surrounding soil lacks sufficient diversity to supply competitive colonisers, the seed community defaults to transmitting what it already contains. This distinction between active host-mediated retention and passive inheritance by coloniser default is important for interpretation and cannot be fully resolved with our current design; an experiment that manipulates soil inoculum diversity while holding treatment chemistry constant would be required to disentangle the two mechanisms. What the contrast between b and c assembly dynamics clearly demonstrates is that soil community richness, itself a product of agricultural management history, exerts a direct and quantifiable influence on how much of each new generation’s seed microbiome is assembled from the environment versus inherited from the maternal lineage.

### MAG-level evidence for persistence and turnover

At the resolution of MAGs, the community dynamics described above are broadly consistent with the genus-level profiles. Metagenomic assembly and binning yielded a total of 485 MAGs, of which 37 were classified as high quality, 75 as medium quality, and 373 as low quality according to MIMAG standards (Table S3). Only the 112 high- and medium-quality MAGs were retained for downstream analyses to ensure taxonomic assignments were based on high-confidence genomic reconstructions. The UpSet plot of genus-level MAG co-occurrence across samples (Figure 5) highlights two results of particular interest. First, sample b contained by far the largest and most exclusive MAG pool in the dataset, with the majority of b-soil MAGs having no representation in any seed sample, consistent with the abundance-without-colonisation pattern described for *Streptomyces* and *Bradyrhizobium* at the genus level, and reinforcing the view that soil richness does not translate proportionally into seed recruitment. Second, the only MAGs shared exclusively between the two Stage 0 seed populations (MODg0 and LANDg0) across both varieties corresponded to *Burkholderia*-related lineages, the same genus identified as displaced after g1 in the abundance profiles. This cross-variety consistency at the MAG level suggests that *Burkholderia* was a detectable component of the maternal seed legacy, providing support for the idea that its subsequent reduction may reflect a genuine community transition rather than a mere sampling artifact. Finally, no MAG-level genera were exclusive to a single treatment lineage across both g1 and g2. The persistent core defined at the genus level therefore did not resolve into stable lineage-specific genomic populations at this sequencing depth. Detecting such signatures would require deeper MAG recovery across replicates ^49,50^.

A notable finding preceding the functional analysis was the complete absence of recoverable high- or medium-quality MAGs from the chemical carbon source soil, where only low-quality bins were retrieved. This suggests that synthetic fertilisation could reduce soil microbial diversity below the threshold required for metagenomic binning at the sequencing depth employed, and is consistent with the lower richness documented in c soils above. As a consequence, no soil-level genomic context is available for the chemical treatment lineage, and this asymmetry should be borne in mind when interpreting these results.

### Plant growth-promoting genes presence across MAGs

To assess whether the persistent and treatment-exclusive genera identified above carry genomic potential for plant growth promotion, we screened all high- and medium-quality MAGs for the presence of fifteen PGP-associated gene categories spanning nitrogen fixation cluster assembly (nifS, nifP, nifU, nifJ, nifV), phosphate solubilisation via the pyrroloquinoline quinone (PQQ) pathway (pqqC, pqqA, pqqD), auxin biosynthesis (iaaH, ipdC, nitrilase), and other PGP-relevant functions including phytase activity (phy), ACC deaminase (acdS), and nitrite/nitrate-related reductases (nrfB/nthB) (Figure 6).

**Figure 5:**
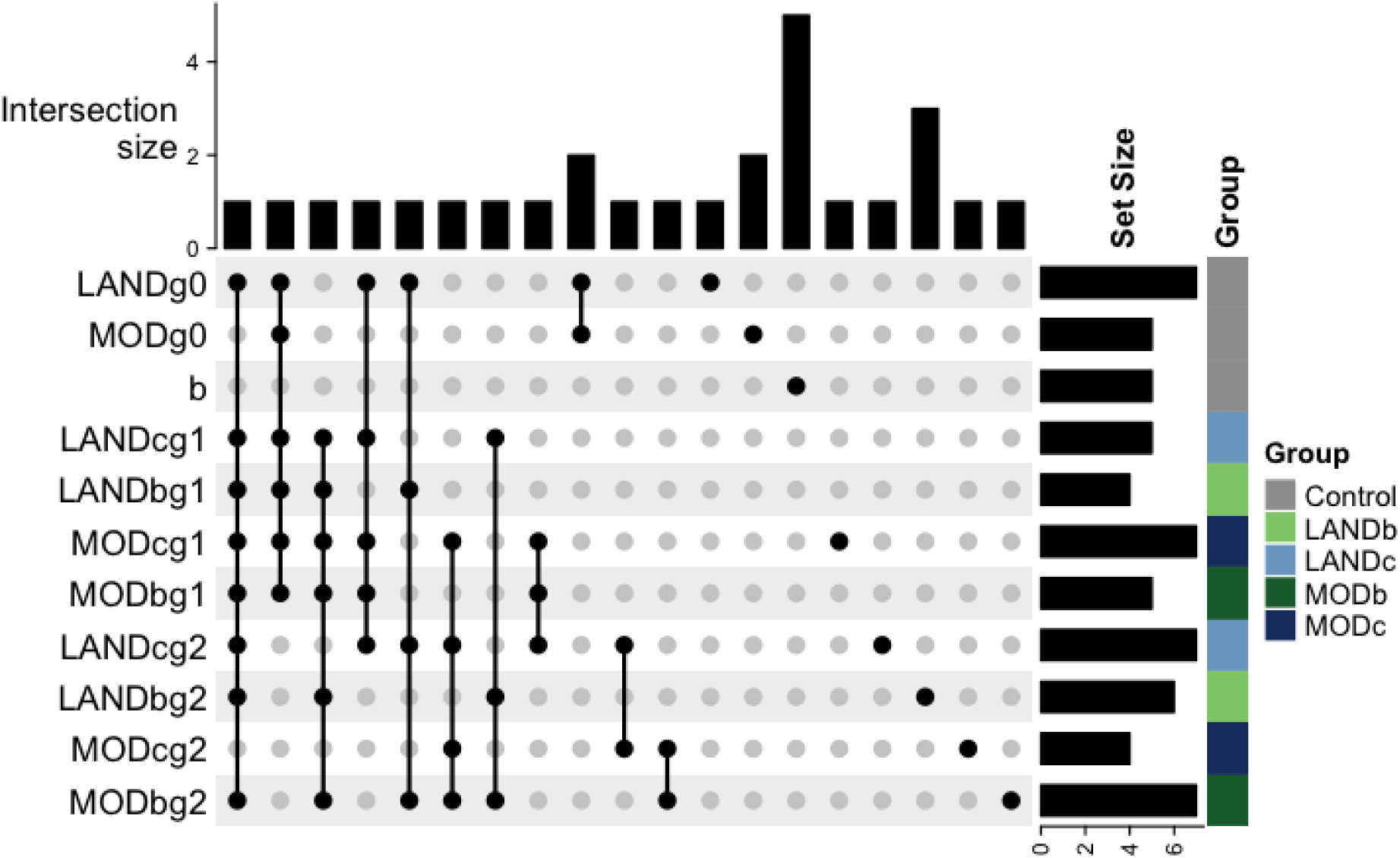
Distribution of unique genera of high- and medium-quality MAGs. Bars above the dot matrix indicate the number of genera shared exclusively across the combination of samples indicated by connected dots; bars annotation on the left indicate total genus-level set size per sample. Soil sample (b) and Stage 0 seeds (MODg0, LANDg0) are annotated in grey, representing environmental baselines without treatment-generation identity. No medium- or high-quality MAGs were recovered from the chemical carbon source soil (c), Treatment-generation seed samples are coloured by variety and carbon source: MOD samples in dark tones (navy for c, dark green for b) and LAND samples in light tones (light blue for c, light green for b). Low-quality MAGs were excluded prior to this analysis.

**Figure 6:**
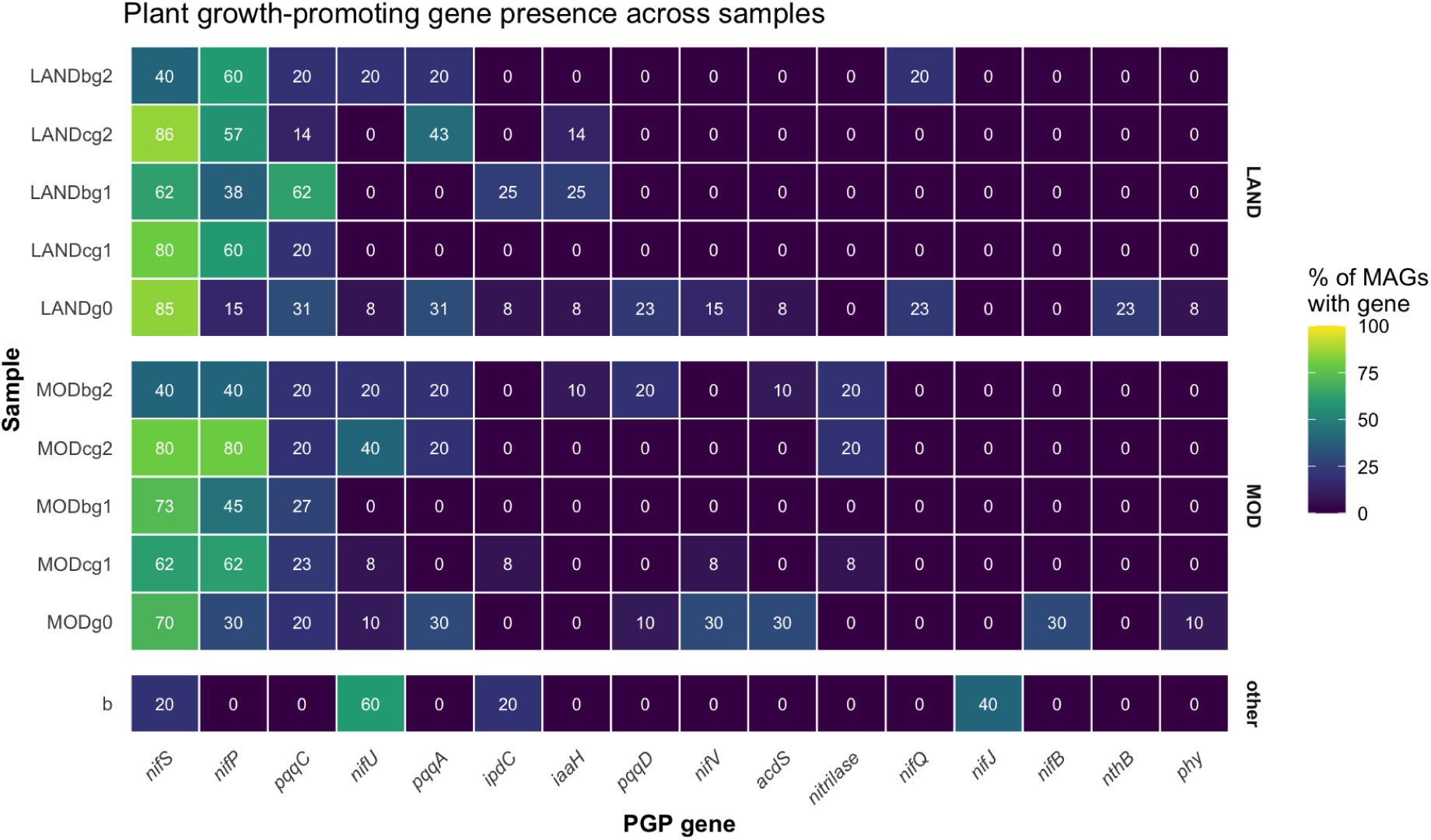
Percentage of MAGs per sample containing direct plant growth promotion genes. The biological carbon source soil (b) is shown in the lower panel; the chemical carbon source soil (c) is absent as no MAGs were recovered from it.

The most broadly distributed genres were nifS, nifP, and pqqC, present at moderate-to-high frequency across nearly all seed treatment groups. nifS encodes a cysteine desulfurase central to iron-sulfur cluster biosynthesis, a process required for nitrogenase maturation but also for a wide range of essential housekeeping enzymes unrelated to nitrogen fixation ^51,52^. Its broad distribution should therefore be interpreted with some caution: the presence of nifS alone is not necessarily diagnostic of active or even potential diazotrophy, because Fe-S cluster assembly genes of this family are found in many genomes irrespective of nitrogen-fixing capacity. By contrast, pqqC is more directly informative of a specific PGP trait. pqqC encodes pyrroloquinoline quinone synthase, the terminal enzyme in PQQ cofactor biosynthesis; PQQ serves as the redox cofactor for membrane-bound quinoprotein glucose dehydrogenase, the enzyme responsible for oxidising glucose to gluconic acid in the periplasm. Gluconic acid production is the principal mechanism by which many soil and rhizosphere bacteria solubilise mineral-bound phosphate, making pqqC presence a reasonably direct proxy for phosphate-solubilising potential ^53^. Its near-ubiquitous presence across seed treatment lineages, peaking at 62% in LANDbg1, suggests that phosphate-acquisition potential is a comparatively conserved trait of the persistent seed-borne community.

Among the more restricted genes, iaaH, nitrilase and ipdC are both associated with bacterial auxin (indole-3-acetic acid, IAA) biosynthesis, acting through different routes: ipdC encodes indole-3-pyruvate decarboxylase, the rate-limiting enzyme of the indole-3-pyruvate pathway, the most common bacterial IAA biosynthesis route and one strongly associated with PGPR-mediated root growth stimulation and lateral root proliferation ^54^; iaaH encodes indoleacetamide hydrolase, part of the alternative indole-3-acetamide pathway more commonly associated with phytopathogenic and some endophytic lineages; and nitrilase catalyzes the direct conversion of indole-3-acetonitrile into IAA. Additionally, acdS encodes 1-aminocyclopropane-1-carboxylate (ACC) deaminase, which degrades the immediate precursor of the plant stress hormone ethylene. This function has well-established links to alleviating root growth inhibition under abiotic stress ^55^. In our dataset, acdS was detected in MAGs from the MOD lineage at Stage 0 and in later generations under both treatments, with more limited detection in LANDg0. Linking these gene detections to specific persistent or displaced genera was not possible at this resolution, as the MAGs used here were assembled independently per replicate rather than from concatenated replicate reads, which affects which genomic bins are recovered and precludes consistent MAG-level tracking of specific taxa across the dataset for this analysis.

The b soil MAG pool (n = 14, Table S3) carried only four of the fifteen screened gene categories (nifS (20%), nifU (60%), ipdC (20%), and nifJ (40%)), and initially appeared to lack pqqC entirely, despite its near-ubiquitous presence across seed lineages. However, subsequent profiling of the unbinned metagenomic assemblies revealed that pqqC is indeed present in the soil community although it evaded MAG recovery. Given that this gene is the most functionally interpretable gene in this screen with respect to phosphate solubilisation, its consistent detection in the seed microbiome highlights its functional priority. In this scenery, phosphate-acquisition potential represents a core trait that is recruited and retained by the seed-associated community across generations.

### Genomic evidence for sporulation potential in the persistent Firmicutes core

To determine whether the persistence of the Firmicutes core across generations is genuinely attributable to spore formation, rather than inferred from phylum identity alone, we screened all high- and medium-quality MAGs for the presence of eight core sporulation genes spanning the master regulator of sporulation onset (spo0A), the sporulation-specific sigma factors that direct forespore- and mother-cell-specific gene expression (sigF, sigE, sigG, sigK), the asymmetric division gene spoIIE, and the SpoIIA operon components spoIIAA and spoIIAB (Figure 7). This marker set was selected based on Galperin et al.’s comprehensive comparative genomic survey of sporulation gene conservation across 180 Firmicutes genomes, which identified this core as the most consistently conserved across spore-forming lineages regardless of taxonomic family, including Bacillaceae and Paenibacillaceae, the families to which the persistent genera identified in this study belong^32^. spo0A is present in essentially every confirmed sporeformer in that survey, and its absence is a reliable negative predictor of sporulation capacity, while the four sigma factors are similarly near-universal among spore-forming taxa regardless of lineage.

**Figure 7:**
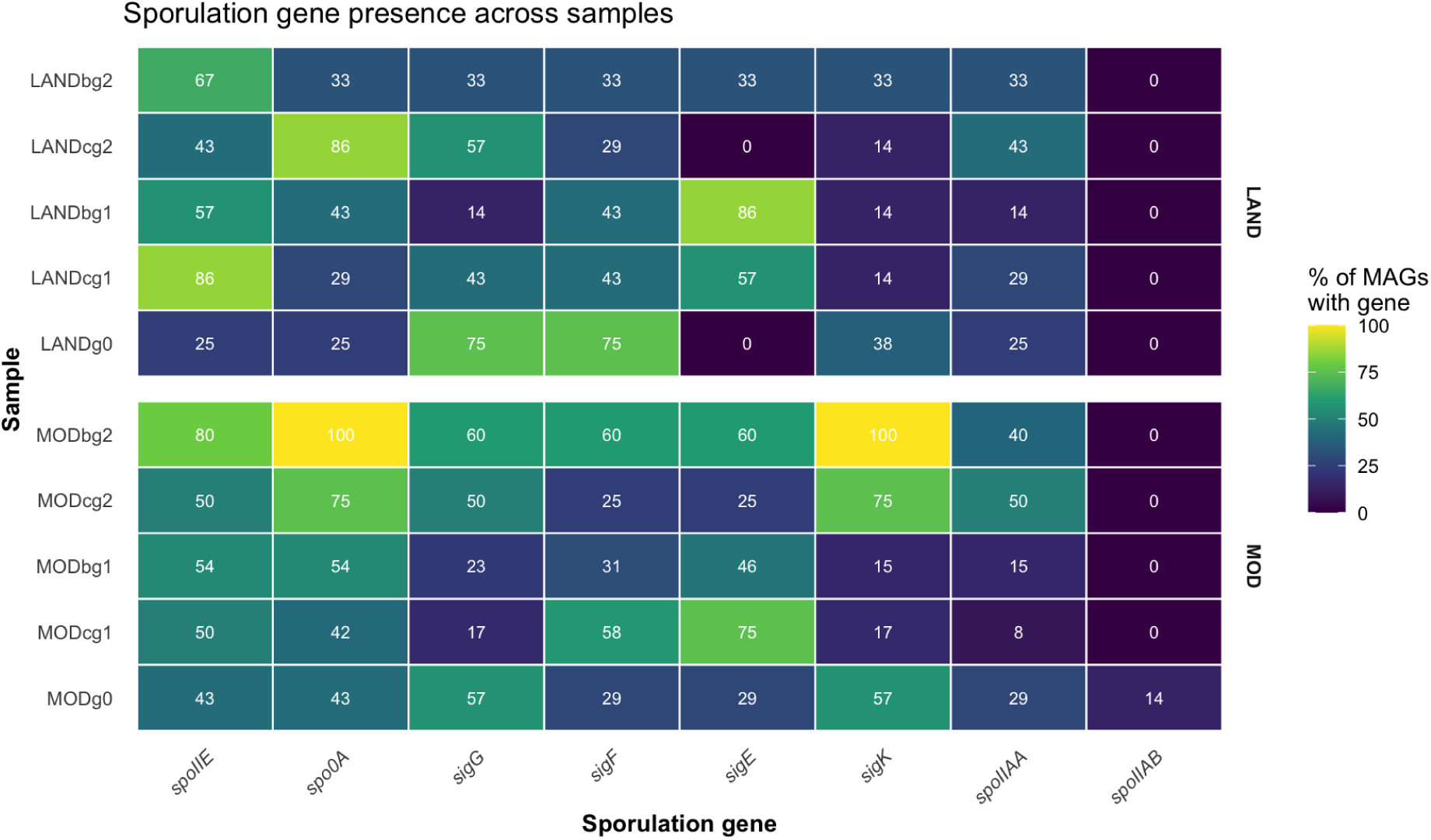
Percentage of high- and medium-quality MAGs per sample group containing core sporulation genes. Marker genes were selected based on their conservation across diverse spore-forming Firmicutes lineages, including Bacillaceae and Paenibacillaceae ^32^.

spo0A and the sigma factors sigF, sigE, sigG, and sigK were detected across nearly every treatment group in both varieties, at frequencies ranging from moderate to high (reaching 100% of MODbg2 MAGs for spo0A and sigK), with spoIIE showing a similarly broad distribution. This pattern provides direct genomic support for sporulation potential within the persistent core MAGs, rather than an inference drawn solely from taxonomic identity. By contrast, spoIIAB was essentially absent across the entire dataset, detected in only a single treatment group (MODg0, 14%), despite the near-universal presence of its operon partner spoIIAA. Given that spoIIAA and spoIIAB are co-transcribed under Spo0A control in the canonical sporulation cascade, this asymmetry may reflect incomplete MAG assembly at this specific locus, lineage-specific streamlining of the SpoIIA regulatory step, a genuine functional substitution not captured by this marker, or even low coverage in that specific region due to the size of the gene; it is presented here without further interpretation, pending closer inspection of the specific MAGs involved. Notably, none of the eight sporulation markers were detected in the biological soil MAG pool (n = 14), and no MAGs were available from the chemical soil for comparison. While the known high prevalence of spore-forming Firmicutis in soil ecosystems initially suggested this might be an artifact of limited MAG recovery, subsequent profiling of the unbinned metagenomic assemblies confirmed the complete absence of these sporulation genes in soil. This indicates a genuine biological feature of the sampled communities, with sporulating bacteria being exclusively inherited.

### Soil community composition as a predictor of seed microbiome assembly

To formally assess whether soil community composition predicted the degree of soil-to-seed recruitment, we used beta regression to explore whether the relative abundance of bacterial classes in soil was associated with the proportion of soil-attributed genera in seed-derived communities, following the approach of Walsh, et al. (2021) ^14^. Eleven classes were significant predictors after Benjamini-Hochberg FDR correction (padj < 0.05). Models were fitted across all replicates of the four soil samples (bg1, bg2, cg1, cg2); given this limited number of observations, these relationships are exploratory and indicate the direction of association rather than providing formal hypothesis tests. Nine of these — Ardenticatenia, Bacteroidia, Cyanophyceae, Spirochaetia, Desulfuromonadia, Rubrobacteria, Nitrospiria, Sphingobacteriia, and Chitinophagia — were present at negligible proportions across all soil samples, only appearing in three or fewer replicates, with predictor ranges spanning less than 0.03. Slopes estimated over such a narrow range are unstable, as small absolute differences are amplified into steep apparent relationships. These nine classes were therefore excluded and are not interpreted further.

The two classes retained in the final model, spanning a biologically informative range of soil proportions, were Alphaproteobacteria and Gammaproteobacteria (Figure S2). These showed opposite directional relationships with soil-to-seed transfer. Alphaproteobacteria were negatively associated with the proportion of soil-derived genera in the seed community (pseudo-R² = 0.44, padj = 0.028): soils richer in Alphaproteobacterial taxa, particularly *Bradyrhizobium*, are associated with seeds with lower proportions of soil-recruited genera. This is ecologically coherent given that *Bradyrhizobium* and its Alphaproteobacterial relatives, despite their dominance in b soils, were undetected across all seed samples and generations. When taxa incapable of seed colonisation dominate the soil pool, they effectively occupy the soil pool at the expense of taxa capable of seed colonisation that could successfully recruit into the seed, suppressing overall soil-to-seed transfer even as total soil diversity increases. Gammaproteobacteria, by contrast, showed a positive relationship with soil-to-seed transfer (pseudo-R² = 0.49, padj = 0.016): soils richer in Gammaproteobacteria were associated with a higher proportion of soil-derived genera in seeds. This is consistent with the known seed-colonisation competence of several Gammaproteobacterial genera, including *Pantoea* and *Pseudomonas*, both recovered in seed samples in our dataset, suggesting that this class contains taxa genuinely equipped to traverse the soil-seed barrier.

Taken together, these two relationships point to a functionally meaningful distinction within the soil community between taxa with and without seed-colonisation competence, and suggest that the class-level composition of the soil, not merely its diversity, determines how much environmental recruitment contributes to seed microbiome assembly. This finding is directly complementary to that of Walsh et al., who reported the reciprocal relationship: higher Gammaproteobacteria abundance in soil inocula was negatively associated with the retention of seed-associated taxa in seedling microbiomes ^14^. As soil-derived and seed-derived fractions are complementary by construction, greater Gammaproteobacterial recruitment into the seed necessarily reduces the retained maternal fraction, so the two results describe the same phenomenon from opposite sides.

## CONCLUSIONS

The maize seed-borne microbiome exhibited both intergenerational persistence and taxonomic turnover under contrasting fertilization regimes. Some taxa, principally spore-forming Firmicutes, demonstrate a clear capacity to persist across generations independent of soil recruitment, while others, such as *Burkholderia*, decline to negligible abundance despite consistent presence in the prior generation and continued availability in the surrounding environment. Under biological farming conditions, roughly half of each generation’s seed microbiome was assembled from the soil and about a third inherited vertically, with the remainder unattributed; under chemical fertilisation, this balance shifted markedly toward vertical transmission. These compositional differences extended to the genomic level: PGP-relevant genes, including pqqC, were most prevalent among seed-associated MAGs under biological treatment and were not detected in the soil MAG pool, being only present in low abundance at reads level, indicating that the functional potential of the seed microbiome is not a direct and proportional subset of what the surrounding soil offers. Whether these transgenerational patterns translate to measurable consequences for plant fitness or persist under the additional stochasticity of field conditions remains to be determined. What the data clearly support is that farming practice does not merely affect the plant that grows in the soil; it shapes the microbial legacy that the plant passes to its offspring. Establishing the mechanisms underlying these patterns, their consequences for plant performance and their persistence under field conditions will require further investigation.

## DATA AVAILABILITY STATEMENT

The raw sequencing data generated in this study have been deposited in the NCBI Sequence Read Archive (SRA) and are available under BioProject accession PRJNA1544141(https://www.ncbi.nlm.nih.gov/bioproject/1544141).

## ACKNOWLEDGEMENTS

During the preparation of this work, the authors used Claude (version Sonnet 4.6, Anthropic) to improve the language, readability, and flow of the manuscript. After using this tool, the authors reviewed and edited the content as needed and take full responsibility for the final publication. This work was supported by Fundação Carlos Chagas Filho de Amparo à Pesquisa do Estado do Rio de Janeiro (FAPERJ; grants E-26/201.117/2022, E-26/200.342/2026, and E-26/210.291/2021), Coordenação de Aperfeiçoamento de Pessoal de Nível Superior (CAPES, Finance Code 001), Conselho Nacional de Desenvolvimento Científico e Tecnológico (CNPq; grants 440221/2022-6, 313024/2023-5, and 420095/2023-3), and Programa de Apoio à Pesquisa, Inovação e Cultura (PAPIC/UENF, SEI-260002/007031/2024). The funding agencies had no role in the design of the study and collection, analysis and interpretation of data and in writing.

## AUTHOR CONTRIBUTIONS

Conceptualization: SHJ, FLO, RSE, TMV Methodology: SHJ, SS, JPM, RCR, GPM, BCR, VJL, UAO, Validation: SHJ, SS, GPM, BCR, VJL, Formal analysis: SHJ, SS, BCR, VJL Investigation: SHJ, TMV Resources: SHK, RNF, FLO Data Curation: SHJ Writing - Original Draft: SHJ Writing - Review and Editing: SHJ, RNF, FLO, RSE, TMV Supervision: RSE, TMV Project administration: FLO, TMV Funding Acquisition: RNF, FLO, TMV

## REFERENCES

1. Hu, H. et al. Seed-Borne Endophytes and Their Host Effects. Microorganisms 13, (2025).

2. Qi, X., Jin, D., Olimi, E., Chen, X. & Cernava, T. What do we know about the seed microbiome? Microbiome 14, (2026).

3. Liu, F. et al. Soil indigenous microbiome and plant genotypes cooperatively modify soybean rhizosphere microbiome assembly. BMC Microbiol 19, 201 (2019).

4. Nelson, E. B., Simoneau, P., Barret, M., Mitter, B. & Compant, S. Editorial special issue: the soil, the seed, the microbes and the plant. Plant Soil 422, 1–5 (2018).

5. Abdelfattah, A., Tack, A. J. M., Lobato, C., Wassermann, B. & Berg, G. From seed to seed: the role of microbial inheritance in the assembly of the plant microbiome. Trends Microbiol 31, 346–355 (2023).

6. Figueiredo Dos Santos, L., et al. Insights into the structure and role of seed-borne bacteriome during maize germination. FEMS Microbiol Ecol 97, (2021).

7. Truyens, S., Weyens, N., Cuypers, A. & Vangronsveld, J. Bacterial seed endophytes: genera, vertical transmission and interaction with plants. Environ. Microbiol. Rep. 7, 40–50 (2015).

8. Henaut-Jacobs, S., Cyríaco, B. E. B., Pedrosa-Silva, F., Olivares, F. L. & Venancio, T. M. Comparative metagenomic profiling of seed-borne microbiomes in a landrace and a hybrid maize variety. Seeds 3, 528–539 (2024).

9. Leibold, M. A. et al. The metacommunity concept: a framework for multi-scale community ecology. Ecol. Lett. 7, 601–613 (2004).

10. Nemergut, D. R. et al. Patterns and processes of microbial community assembly. Microbiol Mol Biol Rev 77, 342–356 (2013).

11. Zhou, F. et al. Diversity Shifts in the Root Microbiome of Cucumber Under Different Plant Cultivation Substrates. Front Microbiol 13, 878409 (2022).

12. Mahmoudi, M., Almario, J., Lutap, K., Nieselt, K. & Kemen, E. Microbial communities living inside plant leaves or on the leaf surface are differently shaped by environmental cues. ISME Commun 4, ycae103 (2024).

13. Costa, O. Y. A., Pijl, A., Houbraken, J., van Lith, W. & Kuramae, E. E. Soil substrate source drives the microbes involved in the degradation of gelatin used as a biostimulant. Appl. Soil Ecol. 189 (2023).

14. Walsh, C. M., Becker-Uncapher, I., Carlson, M. & Fierer, N. Variable influences of soil and seed-associated bacterial communities on the assembly of seedling microbiomes. ISME J 15, 2748–2762 (2021).

15. Ewald, P. W. Transmission modes and evolution of the parasitism-mutualism continuum. Ann N Y Acad Sci 503, 295–306 (1987).

16. Roughgarden, J. Holobiont evolution: Mathematical model with vertical vs. Horizontal microbiome transmission. Philos. Theory Pr. Biol. 12, (2020).

17. Gessner, M. O. et al. Diversity meets decomposition. Trends Ecol Evol 25, 372–380 (2010).

18. Wang, Y., Li, Q. & Li, C. Organic fertilizer has a greater effect on soil microbial community structure and carbon and nitrogen mineralization than planting pattern in rainfed farmland of the Loess Plateau. *Front*. Environ. Sci. 11, (2023).

19. Morris, J. J., Lenski, R. E. & Zinser, E. R. The Black Queen Hypothesis: evolution of dependencies through adaptive gene loss. mBio 3, (2012).

20. Xu, Q., et al. Long-Term Chemical-Only Fertilization Induces a Diversity Decline and Deep Selection on the Soil Bacteria. mSystems 5, (2020).

21. da Costa, P. B. et al. The effects of different fertilization conditions on bacterial plant growth promoting traits: guidelines for directed bacterial prospection and testing. Plant Soil 368, 267–280 (2013).

22. De Coster, W., D’Hert, S., Schultz, D. T., Cruts, M. & Van Broeckhoven, C. NanoPack: visualizing and processing long-read sequencing data. Bioinformatics 34, 2666–2669 (2018).

23. Constantinides, B., Hunt, M. & Crook, D. W. Hostile: accurate decontamination of microbial host sequences. Bioinformatics 39, (2023).

24. Li, H. Minimap2: pairwise alignment for nucleotide sequences. Bioinformatics 34, 3094–3100 (2018).

25. Kolmogorov, M., Yuan, J., Lin, Y. & Pevzner, P. A. Assembly of long, error-prone reads using repeat graphs. Nat Biotechnol 37, 540–546 (2019).

26. Kang, D. D. et al. MetaBAT 2: an adaptive binning algorithm for robust and efficient genome reconstruction from metagenome assemblies. PeerJ 7, e7359 (2019).

27. Chklovski, A., Parks, D. H., Woodcroft, B. J. & Tyson, G. W. CheckM2: a rapid, scalable and accurate tool for assessing microbial genome quality using machine learning. Nat Methods 20, 1203–1212 (2023).

28. Bowers, R. M. et al. Minimum information about a single amplified genome (MISAG) and a metagenome-assembled genome (MIMAG) of bacteria and archaea. Nat Biotechnol 35, 725–731 (2017).

29. Chaumeil, P.-A., Mussig, A. J., Hugenholtz, P. & Parks, D. H. GTDB-Tk: a toolkit to classify genomes with the Genome Taxonomy Database. Bioinformatics 36, 1925–1927 (2019).

30. Wood, D. E., Lu, J. & Langmead, B. Improved metagenomic analysis with Kraken 2. Genome Biol 20, 257 (2019).

31. Henaut-Jacobs, S. et al. Genomic plasticity of the *Azospirillum* genus in a biotechnological context. bioRxiv (2025) doi:10.1101/2025.10.16.682878.

32. Galperin, M. Y., Yutin, N., Wolf, Y. I., Vera Alvarez, R. & Koonin, E. V. Conservation and Evolution of the Sporulation Gene Set in Diverse Members of the *Firimicutes*. J Bacteriol 204, e0007922 (2022).

33. Zhang, C. & Freddolino, L. A large-scale assessment of sequence database search tools for homology-based protein function prediction. Brief Bioinform 25, (2024).

34. Oksanen, J., et al. Vegan: Community ecology package. CRAN: Contributed Packages The R Foundation 10.32614/cran.package.vegan (2001).

35. Paradis, E., et al. Ape: Analyses of phylogenetics and evolution. CRAN: Contributed Packages The R Foundation 10.32614/cran.package.ape (2002).

36. Zeileis, A., Cribari-Neto, F., Grün, B. & Kosmidis, I. betareg: Beta Regression. CRAN: Contributed Packages The R Foundation 10.32614/cran.package.betareg (2004).

37. Compant, S., Clément, C. & Sessitsch, A. Plant growth-promoting bacteria in the rhizo- and endosphere of plants: Their role, colonization, mechanisms involved and prospects for utilization. Soil Biol. Biochem. 42, 669–678 (2010).

38. Hardoim, P. R. et al. The Hidden World within Plants: Ecological and Evolutionary Considerations for Defining Functioning of Microbial Endophytes. Microbiol Mol Biol Rev 79, 293–320 (2015).

39. Fukami, T. Historical contingency in community assembly: Integrating niches, species pools, and priority effects. Annu. Rev. Ecol. Evol. Syst. 46, 1–23 (2015).

40. Sprockett, D., Fukami, T. & Relman, D. A. Role of priority effects in the early-life assembly of the gut microbiota. Nat Rev Gastroenterol Hepatol 15, 197–205 (2018).

41. Barret, M. et al. Emergence shapes the structure of the seed microbiota. Appl Environ Microbiol 81, 1257–1266 (2015).

42. Rybakova, D. et al. The structure of the Brassica napus seed microbiome is cultivar-dependent and affects the interactions of symbionts and pathogens. Microbiome 5, 104 (2017).

43. Compant, S. et al. Harnessing the plant microbiome for sustainable crop production. Nat Rev Microbiol 23, 9–23 (2025).

44. Borriss, R. Bacillus, A plant-beneficial bacterium. in *Principles of Plant-Microbe Interactions 379–391* (Springer International Publishing, Cham, 2015).

45. Perin, L. et al. Diazotrophic burkholderia species associated with field-grown maize and sugarcane. Appl Environ Microbiol 72, 3103–3110 (2006).

46. Compant, S., Nowak, J., Coenye, T., Clément, C. & Ait Barka, E. Diversity and occurrence of Burkholderia spp. in the natural environment. FEMS Microbiol Rev 32, 607–626 (2008).

47. Kumar, A., Longmei, N., Kumar, P. & Kaushik, P. Molecular marker analysis of genetic diversity in maize: A review. OBM Genet. 06, 1–19 (2022).

48. Santiago, Malvar, R. A., Revilla, P. & Butrón, A. Maize landraces as useful donors of genetic diversity for resilience to drought. J. Agric. Food Res. 23, 102297 (2025).

49. Orellana, L. H., Krüger, K., Sidhu, C. & Amann, R. Comparing genomes recovered from time-series metagenomes using long- and short-read sequencing technologies. Microbiome 11, 105 (2023).

50. Rocha, U. et al. Simulation of 69 microbial communities indicates sequencing depth and false positives are major drivers of bias in prokaryotic metagenome-assembled genome recovery. PLoS Comput Biol 20, e1012530 (2024).

51. Olson, J. W., Agar, J. N., Johnson, M. K. & Maier, R. J. Characterization of the NifU and NifS Fe-S cluster formation proteins essential for viability in Helicobacter pylori. Biochemistry 39, 16213–16219 (2000).

52. Tokumoto, U., Kitamura, S., Fukuyama, K. & Takahashi, Y. Interchangeability and distinct properties of bacterial Fe-S cluster assembly systems: functional replacement of the isc and suf operons in Escherichia coli with the nifSU-like operon from Helicobacter pylori. J Biochem 136, 199–209 (2004).

53. Matsushita, K. et al. Escherichia coli is unable to produce pyrroloquinoline quinone (PQQ). Microbiology (Reading) 143 **( Pt** **10**), 3149–3156 (1997).

54. Spaepen, S. & Vanderleyden, J. Auxin and plant-microbe interactions. Cold Spring Harb Perspect Biol 3, (2011).

55. Glick, B. R. Stress Control and ACC Deaminase. in Principles of Plant-Microbe Interactions 257–264 (Springer International Publishing, Cham, 2015).

