## Supplementary material for "Vertical transmission and community turnover in seed-borne microbiomes: lessons from an aMAIZEing model system": Suplementary figures


Figure S1 - Principal component analysis showing the difference in taxa composition across samples (A) separated by individual combinations of treatment and generation, and (B) highlighting contrasting characteristics for genotype and soil separately.

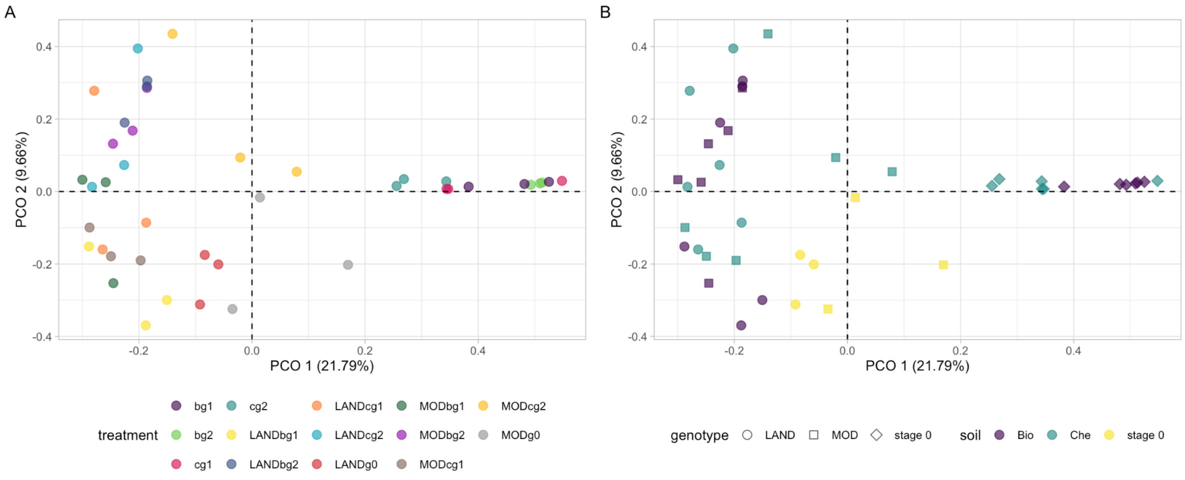


Figure S2 - Beta regression showing the degree of soil-to-seed recruitment. In the Y-axis we show the proportion of bacteria in the sample originally marked as soil-borne, in the X-axis we see the proportion of each class in the soil.

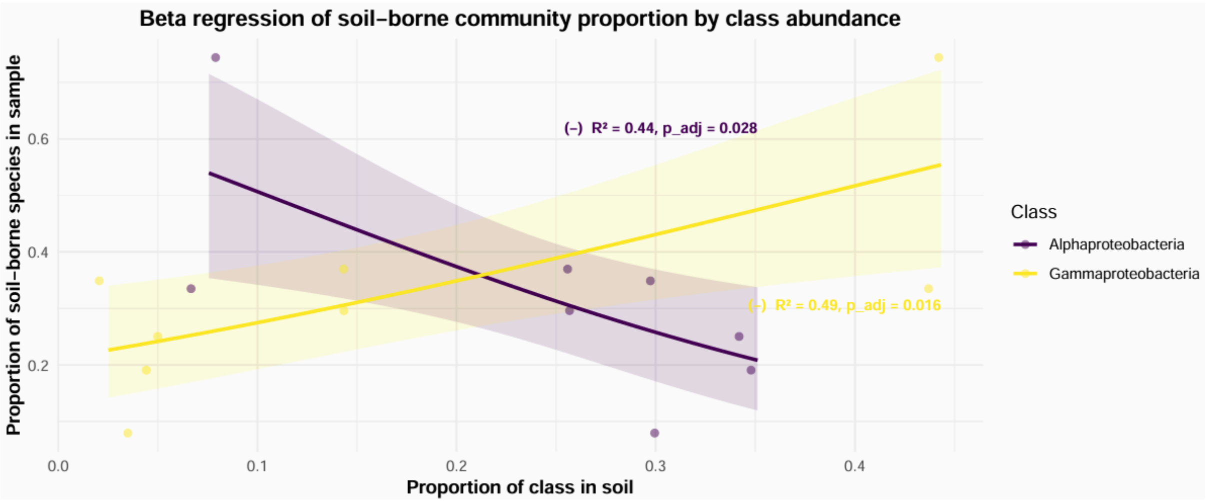
Beta regression of soil–borne community proportion by class abundance
(–) R² = 0.44, p_adj = 0.028
(+) R² = 0.49, p_adj = 0.016
Proportion of soil–borne species in sample
Proportion of class in soil
0.0
0.1
0.2
0.3
0.4
0.2
0.4
0.6
Class
Alphaproteobacteria
Gammaproteobacteria
